# Expression and function of SLC38A5, an amino acid-coupled Na^+^/H^+^ exchanger, in triple-negative breast cancer and its relevance to macropinocytosis

**DOI:** 10.1101/2021.06.17.448844

**Authors:** Sabarish Ramachandran, Souad Sennoune, Monica Sharma, Muthusamy Thangaraju, Varshini Suresh, Yangzom D. Bhutia, Kevin Pruitt, Vadivel Ganapathy

**Affiliations:** Department of Cell Biology and Biochemistry, Texas Tech University Health Sciences Center, Lubbock, TX 79430, USA; Department of Immunology and Molecular Microbiology, Texas Tech University Health Sciences Center, Lubbock, TX 79430, USA; Department of Biochemistry and Molecular Biology, Medical College of Georgia, Augusta University, Augusta, GA 30912, USA; Department of Biology, Texas Tech University, Lubbock, TX 79409, USA

**Keywords:** glutamine addiction, one-carbon metabolism, intracellular alkalinization, breast cancer, macropinocytosis, amiloride

## Abstract

Metabolic reprogramming in cancer cells necessitates increased amino acid uptake, which is accomplished by upregulation of specific amino acid transporters. Since amino acid transporters differ in substrate selectivity, mode of transport, and driving forces, not all tumors rely on any single amino acid transporter for this purpose. Here we report on the differential upregulation of the amino acid transporter SLC38A5 in triple-negative breast cancer (TNBC). The upregulation is evident in primary TNBC tumors, conventional TNBC cell lines, patient-derived xenograft TNBC cell lines, and a mouse model of spontaneous mammary tumor representing TNBC. The upregulation is confirmed by functional assays. SLC38A5 is an amino acid-dependent Na^+^/H^+^ exchanger which transports Na^+^ and amino acids into cells coupled with H^+^ efflux. Since the traditional Na^+^/H^+^ exchanger is an established inducer of macropinocytosis, an endocytic process for cellular uptake of bulk fluid and its components, we examined the impact of SLC38A5 on macropinocytosis in TNBC cells. We found that the transport function of SLC38A5 is coupled to induction of macropinocytosis. Surprisingly, the transport function of SLC38A5 is inhibited by amilorides, the well-known inhibitors of Na^+^/H^+^ exchanger, possibly related to the amino acid-dependent Na^+^/H^+^ exchange function of SLC38A5. The Cancer Genome Atlas database corroborates SLC38A5 upregulation in TNBC. This represents the first report on the selective expression of SLC38A5 in TNBC and its role as an inducer of macropinocytosis, thus revealing a novel, hitherto unsuspected, function for an amino acid transporter that goes beyond amino acid delivery but is still relevant to cancer cell nutrition.

**Summary Statement:** SLC38A5 is an amino acid-coupled Na^+^/H^+^ exchanger that is upregulated in triple-negative breast cancer, and its function in cancer cells goes beyond amino acid delivery; it promotes macropinocytosis, a distinct form of endocytic process for cellular uptake of proteins and other nutrients present in extracellular fluid.

## Introduction

Cancer cells reprogram their metabolism to best suit their need for accelerated proliferation associated with enhanced protein/lipid synthesis, DNA/RNA synthesis, and potentiated machinery for protection against oxidative stress and apoptosis [1–3]. This requires increased supply of nutrients to feed into the altered metabolic pathways; amino acids represent an important group among such nutrients. Because of their hydrophilic nature, amino acids cannot permeate plasma membrane by diffusion; they enter the cells via selective transporters. There are more than three dozen amino acid transporters in mammalian cells [4]. They are grouped into seven different solute carrier gene families, based on the similarity in primary amino acid sequence. The increased demand for amino acids in cancer cells is met by upregulation of selective amino acid transporters. The most studied transporters for their role in cancer include SLC1A5 (Alanine-Serine-Cysteine Transporter 2 or ASCT2) [5–8], SLC7A5 (system L Amino acid Transporter 1 or LAT1) [5–9], SLC7A11 (system x^-^_c_ transporter or xCT) [5,6,10,11], and SLC6A14 (Amino acid Transporter B^0,+^ or ATB^0,+^) [5–8,12–14]. The contribution of each of these transporters to tumor promotion is the provision of selective amino acids to cancer cells, which might impact on a multitude of downstream metabolic pathways and signaling cascades.

SLC38 family contains several amino acid transporters [15], but there is little or no information in the literature on the role of these transporters in cancer. Only recently, a couple of studies have focused on SLC38A2 in breast cancer [16,17]. This transporter represents the classical Na^+^-coupled neutral amino acid transporter known as system A, cloned and functionally characterized in our laboratory [18]. It was called “system A” because of its preference for the amino acid alanine. However, it transports not only alanine but also other small, hydrophilic, amino acids such as glycine and serine. Interestingly, glutamine is an excellent substrate for this transporter. SLC38A2 is induced by hypoxia and its expression is related to endocrine resistance in breast cancer [16]. In triple-negative breast cancer (TNBC), the level of SLC38A2 expression is associated with worse prognosis [17]. Both studies have implicated SLC38A2 in supplying glutamine to cancer cells. Notwithstanding these data suggesting a role for this transporter in breast cancer progression, its expression is actually downregulated in breast cancer, particularly in TNBC (*p*<1.5×10^−8^) (TCGA database). The decreased expression in cancer is paradoxical and counterintuitive, given the proposed role of the transporter as a tumor promoter.

In the present study, we focused on another member of the SLC38 family, namely SLC38A5. This transporter was also cloned in our laboratory [19,20]. It is referred to as SN2 transporter, meaning that it is the second isoform with functional features defined selectively for the amino acid transport system N. This system is Na^+^-coupled and accepts as substrates amino acids that contain nitrogen in the side chain (i.e., glutamine, asparagine, and histidine). However, functional characterization of the cloned transporter revealed two interesting features: (i) it also transports glycine, serine and methionine, with highest affinity towards serine, and (ii) it utilizes an outward-directed H^+^ gradient as a driving force in addition to an inward-directed Na^+^ gradient [19,20]. The transport mechanism involves the transfer of Na^+^ and amino acid substrate in one direction coupled to the transfer of H^+^ in the opposite direction, with the overall transport process being electroneutral. Stated differently, SLC38A5 is an amino acid-coupled Na^+^/H^+^ exchanger. The rationale for the present study to evaluate the expression and function of SLC38A5 in breast cancer is the following: (i) SLC38A5 is a target for the oncogene c-Myc [21]; (ii) the substrate selectivity of SLC38A5 suggests a potentially a key role for this transporter in cancer-associated metabolic pathways such as glutaminolysis (glutamine) and one-carbon metabolism (serine, glycine, and methionine) [14]; (iii) the efflux of H^+^ coupled to amino acid entry fulfills dual needs in cancer cells, namely supply of amino acids and removal of H^+^ [19,20], and (iv) Na^+^/H^+^ exchanger is known to promote macropinocytosis, a novel mechanism for cellular entry of nutrients from extracellular medium [22,23], suggesting a possibility that SLC38A5 as an amino acid-coupled Na^+^/H^+^ exchanger might also promote macropinocytosis in cancer cells.

## Materials and Methods

### Materials

[2,3-^3^H]-L-Serine (specific radioactivity, >5 Ci/mmol) was purchased from Moravek, Inc. (Brea, CA, USA). [2-^3^H]-Glycine (specific radioactivity, x Ci/mmol) and [3,4-^3^H]-glutamine (specific radioactivity, x Ci/mmol) were purchased from Perkin Elmer Corp. (Waltham, MA, USA). Ethylisopropyl amiloride and hexamethylene amiloride were from Sigma Aldrich (St. Louis, MO, USA). Amiloride, benzyl amiloride and harmaline were from Cayman Chemical (Ann Arbor, MI, USA).

### Human tissues

Human breast cancer tissues and the surrounding normal tissues were obtained from the Augusta University Tumor Bank. The Tumor Bank collects and maintains a repository of de-identified tumor tissues and matched normal tissues; the tumor tissue collection has the approval from the Institutional Review Board and the Human Assurance Committee. These tissues are available to investigators without a separate approval from the Institutional Review Board.

### Animals

We used three different transgenic mouse models of spontaneous breast cancer for the analysis of Slc38a5 expression: MMTV-Neu (a model for HER2-positive breast cancer), MMTV-HRAS (a model for Ras activation-associated breast cancer), and MMTV-PyMT (polyoma middle T antigen-driven breast cancer, which starts initially as estrogen receptor positive and subsequently turns into estrogen receptor-negative). These mouse lines were originally obtained from the Jackson Laboratory (Bar Harbor, ME, USA): MMTV-Neu, stock # 002376; MMTV-HRAS, stock # x; MMTV-PyMT, stock # 002374). Since all mice used in the present study never went through pregnancy, we used mammary tissues from adult (~12-week-old) virgin wild type mice as a control for comparison. These mice form spontaneous tumors in mammary glands at different ages: 8-10 months in MMTV-Neu and MMTV-HRAS mouse lines; 2-3 months in the MMTV-PyMT mouse line). The protocol was approved by the Institutional Animal Care and Use Committee of the Texas Tech University Health Sciences Center, Lubbock, TX, USA (IACUC approval number: 15002 for breeding protocol and 18005 for experimental protocol). All the experiments described in this study, including the animal experiments, were conducted at this institution. At the termination of the study, mice were killed by cervical dislocation under CO_2_ anesthesia in accordance with the guidelines from the American Veterinary Medical Association.

### Cell lines and culture conditions

We used four human non-transformed mammary epithelial cell lines and seven estrogen receptor-negative breast cancer cell lines. The non-transformed cell lines are: HMEC (ATCC, Cat. no. PCS-600-010), HBL100 (ATCC, Cat. no. HTB-124), MCF10A (ATCC, Cat. no. CRL-10317), and MCF12A (ATCC, Cat. no. CRL10782). HMEC cells were cultured in Mammary Epithelial Cell Basal Medium (ATCC, Cat. no. PCS-600-030) and HBL100 cells were cultured in modified McCoy’s 5a medium (ATCC, Cat. no. 30-2007). MCF10A and MCF12A cells were cultured in a special medium consisting of Dulbecco’s Modified Eagle’s Medium and Ham’s F12 medium, in a 1:1 ratio, supplemented with 20 ng/ml human EGF, 0.01 mg/ml bovine insulin, 500 ng/ml hydrocortisone, and 100 ng/ml cholera toxin. All media contained 10% fetal bovine serum. Six of the seven estrogen receptor-negative cell lines were obtained from ATCC: BT20 (Cat. no. HTB-19), HCC1937 (Cat. no. CRL-2336), MDA-MB231 (Cat. no. CRM-HTB-26), MDA-MB436 (Cat. no. HTB-136), MDA-MB453 (Cat. no. HTB-131) and MDA-MB468 (Cat. no. HTB-132). SUM1315MO2 cell line was obtained from Expasy (Cat. no. CVCL_5589). BT20 cells were cultured in Eagle’s Minimum Essential Medium (ATCC, Cat. no. 30-2003); HCC1937 cells were cultured in RPMI-1640 medium (ATCC, Cat. no. 30-2001); MDA-MB436 cells were cultured in Leibovitz’s L-15 medium (ATCC, Cat. no. 30-2008), supplemented with 10 μg/ml insulin and 16 μg/ml glutathione. MDA-MB231, MDA-MB453, and MDA-MB468 cells were cultured in Leibovitz’s L-15 medium. SUMO1315MO2 cells were cultured in Ham’s F12 medium containing 10 ng/ml EGF and 5 μg/ml insulin. All media contained 10% fetal bovine serum. All cell lines were mycoplasma-free. Three patient-derived xenograft cell lines were provided by the TTUHSC Cancer Center; these are identified as TXBR-100, TXBR-237 and TXBR-247. These cell lines were cultured in the special medium described above for MCF10A.

### Uptake assays

Uptake of [^3^H]-serine was used to monitor the transport function of SLC38A5. Since SLC38A5 is a Na^+^-coupled transporter with involvement of H^+^ movement in the opposite direction, the uptake assays were done using a pH 8.5 buffer to create an outwardly directly H^+^ gradient across the plasma membrane. As there are several amino acid transporters, even for serine that is used as the substrate in the present study, that are Na^+^-coupled, we cannot specifically monitor the function of SLC38A5 by using Na^+^-containing uptake buffer. However, unlike other Na^+^-coupled amino acid transporters, SLC38A5 is tolerant to Li^+^, meaning that this transporter functions when Na^+^ is replaced with Li^+^ [19,20]. Therefore, we used an uptake buffer with LiCl in place of NaCl. The composition of the uptake buffer was 25 mM Tris/Hepes, pH 8.5, containing 140 mM LiCl, 5.4 mM KCl, 1.8 mM CaCl_2_, 0.8 mM MgSO_4_, and 5 mM D-glucose. Serine is also a substrate for SLC7A5 (LAT1), which is a Na^+^-independent transporter; therefore, uptake via this transporter will contribute to the total uptake measured in the LiCl-containing buffer. To suppress serine uptake that might occur via SLC7A5, the uptake buffer contained 5 mM tryptophan to compete with and block the transport of serine mediated by SLC7A5; SLC38A5 does not transport tryptophan and therefore SLC38A5-mediated serine uptake is not affected by tryptophan. To determine the contribution of diffusion to the total uptake of serine, the same uptake buffer but with LiCl replaced isosmotically with N-methyl-D-glucamine chloride (NMDGCl) was used. Serine uptake was measured in two buffers: (i) LiCl-buffer, pH 8.5 with 5 mM tryptophan; (ii) NMDGCl-buffer, pH 8.5 with 5 mM tryptophan. The uptake in NMDGCl-buffer was subtracted from the uptake in LiCl-buffer to determine specifically the transport activity of SLC38A5.

Cells were seeded in 24-well culture plates (2 × 10^5^ cells/well) with the culture medium. On the day of uptake measurement, the culture plates were kept in a water bath at 37 °C. The medium was aspirated and the cells were washed with uptake buffers. The uptake medium (250 μl) containing [^3^H]-serine was added to the cells. Following incubation for 15 min, the medium was removed and the cells washed three times with ice-cold uptake buffer. The cells were then lysed in 1% sodium dodecyl sulfate/0.2 N NaOH and used for measurement of radioactivity. In the initial experiments for the identification of the transport function of SLC38A5, [^3^H]-glutamine and [^3^H]-glycine were also used as the substrates. But, for most of the experiments, [^3^H]-serine was used.

### Impact of intracellular acidification on SLC38A5 transport activity

Intracellular acidification in cultured cells was accomplished by the NH_4_Cl pretreatment as described previously [24]. Cells were cultured in 24-well culture plates. On the day of the uptake assay, cells were incubated with NaCl-buffer (same as the uptake buffer except that LiCl or NMDGCl was replaced with NaCl), pH 7.5. containing 25 mM NH_4_Cl for 30 min. Osmolality of the buffer was maintained by adjusting the concentration of NaCl (115 mM instead of 140 mM). Cells were then washed with the same buffer, but in the absence of NH_4_Cl (concentration of NaCl was increased to 140 mM), and then used to monitor the transport activity of SLC38A5 as described above. During the preincubation with NH_4_Cl, intracellular pH increases because of the conversion of NH_3_ to NH_4_^+^ inside the cells and then the pH comes down to physiological pH because of the various pH regulatory mechanisms. During the subsequent wash in the absence of NH_4_Cl, NH_4_^+^ within the cells dissociates to NH_3_ and H^+^, and NH_3_ diffuses out of the cell, leaving H^+^ inside with consequent intracellular acidification.

### RT-PCR

RNA was extracted from cells and mammary tissues from mice (normal and tumor) by TRIzol reagent (Thermo Fisher Scientific), and the RNA was reverse-transcribed using High-capacity cDNA reverse transcription kit (Thermo Fisher Scientific). PCR and quantitative-PCR were performed with Takara Taq Hot Start Version (TaKaRa Biotechnology, Shiga, Japan) or Power SYBR Green PCR master mix (Thermo Fisher Scientific). Human-specific primers were used for cell lines whereas mouse-specific primers were used for mouse tissues. Primer sequences are shown in Supplementary Table 1. The relative mRNA expression was determined by the 2^-ΔΔ^Ct method. HPRT (hypoxanthine/guanine phosphoribosyl transferase) was used as a housekeeping gene for normalization.

### Macropinocytosis assay

Cells were plated on coverslips, placed in wells in a 12 well-plate, at a density of 1×10^5^ cells/well, and cultured with 5% CO_2_ at 37 °C using the culture media (with 10% fetal bovine serum) recommended for the respective cell lines. The cells were allowed to reach ~70% confluency. About 16 h prior to macropinocytosis assay, the medium was removed and fresh culture medium without the fetal bovine serum was added. Cells were washed three times with buffers consisting of 140 mM NaCl, LiCl or NMDGCl buffer, pH 7.5, all of them containing the following as the common components: 5.4 mM KCl, 1.8 mM CaCl_2_, 0.8 mM MgSO_4_, and 5 mM D-glucose. Subsequently, the cells were exposed to the same respective buffers, but now containing TMR-dextran (100 μg/ml), with or without amino acids, and in the absence or presence of ethylisopropyl amiloride (EIPA) (100 μM) for 15 min at 37 °C. Then, the cells were washed three times with the respective buffer alone and then fixed with 4% paraformaldehyde for 5 min, washed several times with phosphate-buffered saline, and mounted using Prolong diamond with 4,6-diamidino-2-phnenylindole (DAPI) as a nuclear marker. Cell images were taken using a Nikon confocal microscope, with a 60X objective and analyzed using the Nippon Ichi software. The images represent a maximum projection intensity derived from a Z-stack. The fluorescence quantification was done by measuring the Corrected Total Cell fluorescence (CTCF) using Image J and the following formula:

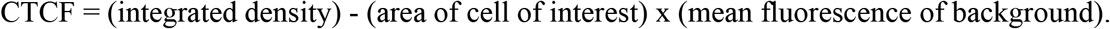

For each cell in an image, an outline was drawn to measure integrated density, area of the cell of interest, and mean fluorescence of the adjacent background around the cell of interest.

### Statistics

RT-PCR and uptake studies were done in triplicates and repeated twice. Statistical analysis was performed with a two-tailed, paired Student’s t-test for single comparison and a *p* value < 0.05 was considered statistically significant. Data are given as means ± S.E. For quantification of fluorescence signals in image analysis related to micropinocytosis and for analysis of inhibition of serine uptake by amiloride and its derivatives, ANOVA followed by Dunn’s test was used to determine the significance of difference among the different groups.

## Results

### Differential expression of SLC6A14 and SLC38A5 in estrogen receptor-positive (ER+) breast cancer and triple-negative (TNBC) breast cancer

We first examined primary tumor specimens from patients with ER+ breast cancer and TNBC for expression of SLC6A14 and SLC38A5. These were the same specimens used in a previously published study to examine the expression of SLC6A14 [25]. As reported earlier [25], SLC6A14 was expressed ER+ breast cancer but not in TNBC (Fig. 1). In contrast, there was a marked upregulation of SLC38A5 in TNBC (~3-fold; p<0.001; n=4). There was a small increase in the expression of this transporter in ER+ breast cancer compared to corresponding adjacent normal tissues, but the difference was not statistically significant. We also studied the expression of SLC1A5. We found the expression of this transporter to be slightly increased in breast cancer but with no differential expression in ER+ breast cancer versus TNBC.

**Fig. 1.**
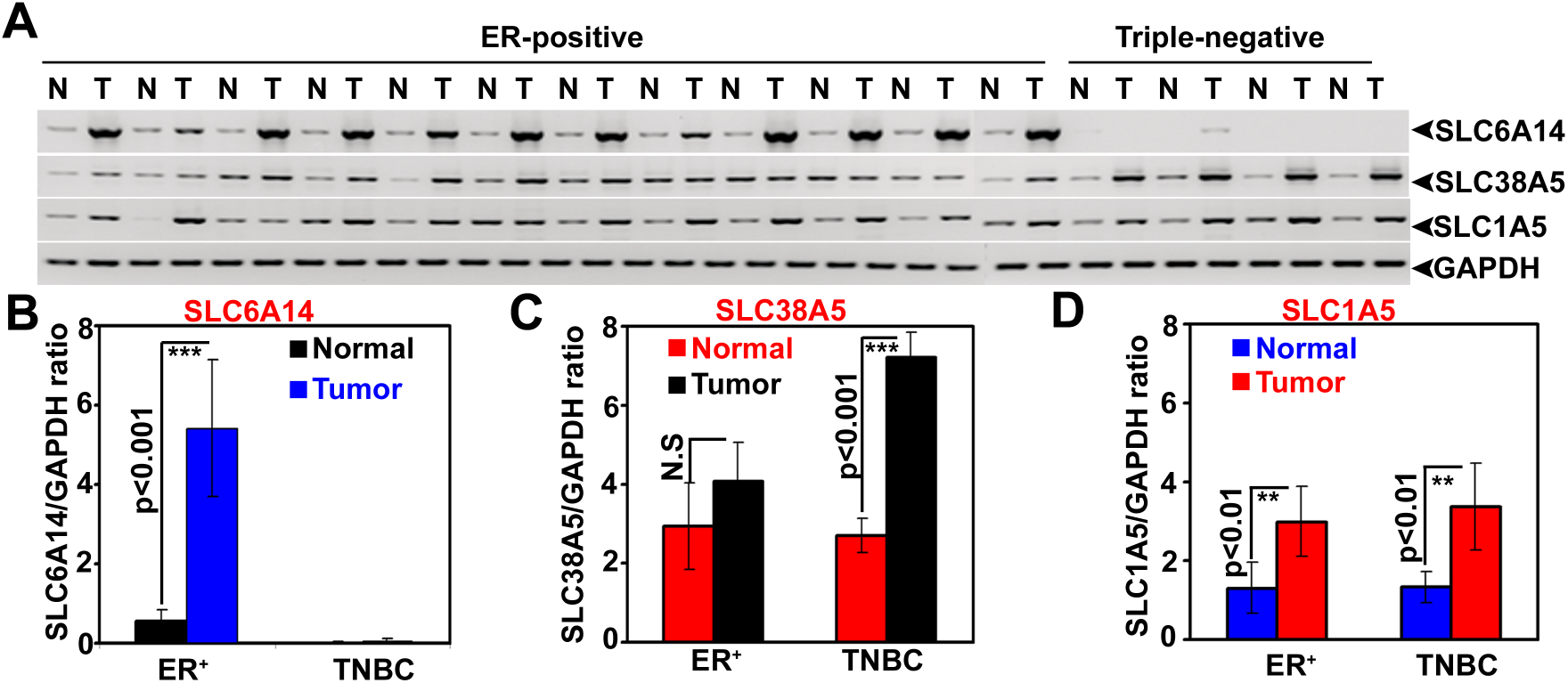
Relative expression of the amino acid transporters SLC6A14, SLC38A5, and SLC1A5 in paired tissues samples of breast cancer (ER+ breast cancer and ER-negative breast cancer) and normal mammary gland. (A) Semi-quantitative RT-PCR (N, normal mammary gland; T, tumor tissue). (B) Quantitative RT-PCR for SLC6A14. (C) Quantitative RT-PCR for SLC38A5. (D) Quantitative RT-PCR for SLC1A5. ER, estrogen receptor; TNBC, triple-negative breast cancer; GAPDH, glyceraldehyde-3-phosphate dehydrogenase.

We then screened TNBC cell lines for expression of various transporters that have glutamine as a common substrate. We focused on the transporters for glutamine because of its multiple biological functions in cancer cell proliferation and growth. For comparison, we used four non-malignant breast epithelial cell lines. Interestingly, the only amino acid transporter that was upregulated in TNBC cell lines was SLC38A5 (Fig. 2). There was no difference in the expression of SLC1A5 and SLC7A5 between non-malignant and TNBC cell lines. We also monitored the expression of two other members in the SLC38 family, namely SLC38A1 and SLC38A2, both of which accept glutamine as a substrate. Two recent studies have reported that SLC38A2 plays a critical role in supplying glutamine to breast cancer cells [16, 17]. But, we did not find any difference in the expression of these two transporters between non-malignant and TNBC cell lines (Fig. 2A).

**Fig. 2.**
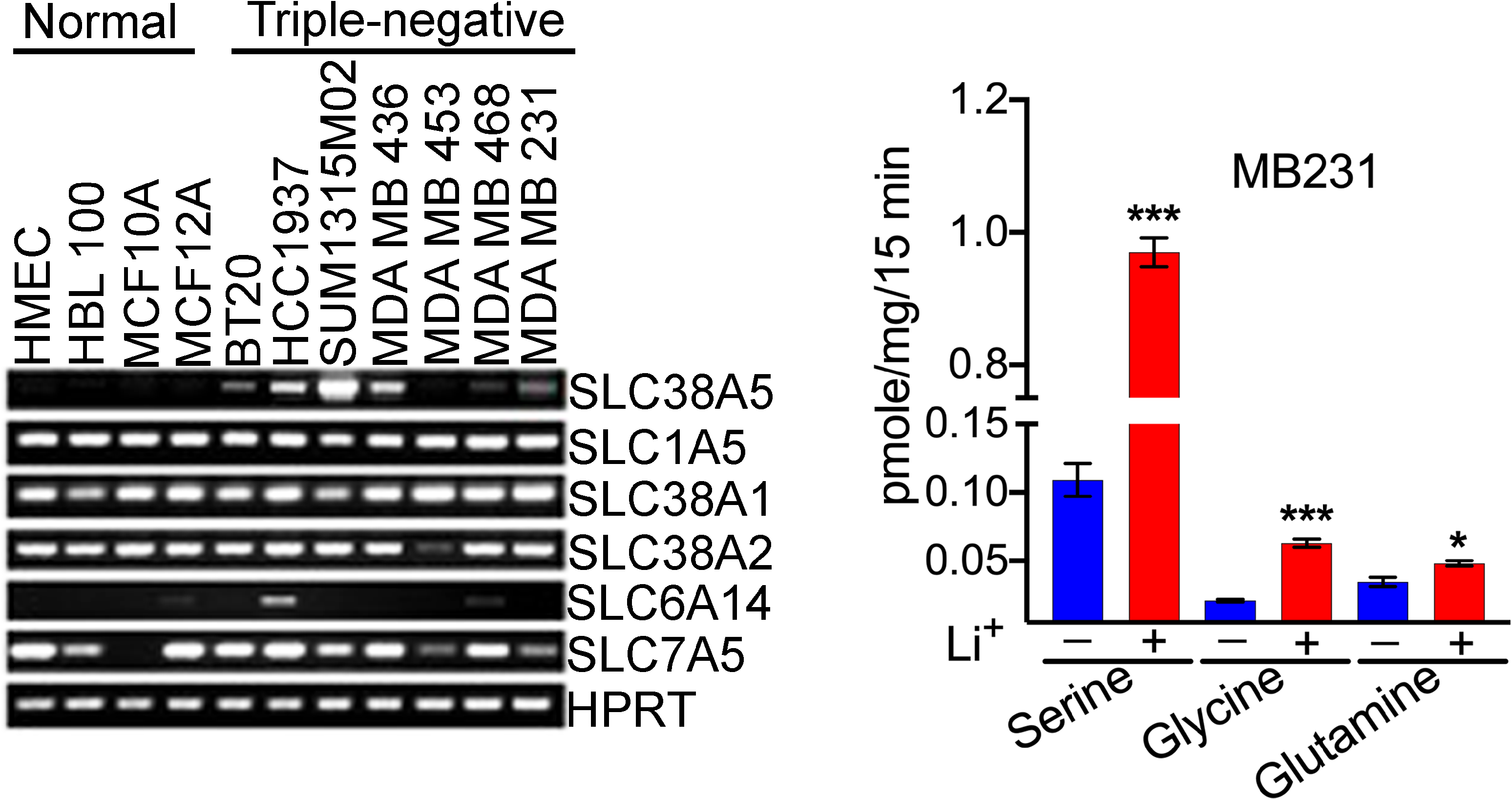
Differential expression of SLC38A5 and SLC6A14 in TNBC cell lines. (A) RT-PCR analysis of mRNAs for different amino acid transporters in four non-malignant breast epithelial cell lines and seven TNBC cell lines. HPRT, hypoxanthine/guanine phosphoribosyl transferase. (B) Transport activity of SLC38A5 in the TNBC cell line MB231 as monitored by the uptake of serine, glycine, and glutamine as the substrates for the transporter. The transport function of SLC38A5 was monitored in an uptake buffer (pH 8.5) containing 5 mM tryptophan to suppress the involvement of SLC7A5 in the uptake and by comparing the uptake in the presence and absence of Li^+^. The Li^+^-stimulatable uptake under these uptake conditions was taken as the transport activity specific for SLC38A5. *, p<0.05; ***, p<0.001, compared to uptake in the absence of Li^+^.

### Functional evidence for SLC38A5 expression in TNBC cells

A unique feature that distinguishes SLC38A5 from other Na^+^-coupled amino acid transporters is its ability to function in the presence of Li^+^ in place of Na^+^. We have confirmed this feature of Li^+^ tolerance using cloned human SLC38A5 [19] and rat SLC38A5 [20]. As SLC38A5 recognizes glutamine, glycine, and serine as substrates [19, 20], we examined the transport activity for these three amino acids in MB231 cells, a widely used prototypical TNBC cell line, in the presence and absence of Li^+^. The uptake buffer (pH 8.5) with Li^+^ was prepared by isosmotically replacing NaCl with LiCl. The buffer without Li^+^ was prepared by isosmotically replacing NaCl with NMDGCl. We found robust uptake of serine in these cells in LiCl-buffer (Fig. 2B). The uptake in the absence of Li^+^ was only 10% of the total uptake measured in the presence of Li^+^. Glutamine uptake and glycine uptake were also increased in the presence of Li^+^, but the fold-stimulation elicited by Li^+^ was smaller compared to what was seen with serine uptake. Serine is the most favored substrate for SLC38A5, and the most prominent Li^+^-dependent uptake for this amino acid confirms this substrate selectivity.

We also used patient-derived xenograft (PDX) TNBC cell lines for the analysis. We obtained three PDX cell lines from the Texas Cancer Cell Repository (www.TXCCR.org), which were developed from PDX samples representing TNBC. First, we monitored the expression of SLC6A14 and SLC38A5 in these cell lines and found all three cell lines to exhibit robust expression of SLC38A5 (Fig. 3A). One of the three cell lines was positive for SLC6A14. We then selected one of the cell lines to monitor the transport function for SLC38A5. Li^+^-dependent serine uptake was robust in the TXBR-100 cell line (~10-fold stimulation by Li^+^) (Fig. 3B). Glycine uptake and glutamine uptake were also stimulated by Li^+^, but the fold-stimulation was the best for serine.

**Fig. 3.**
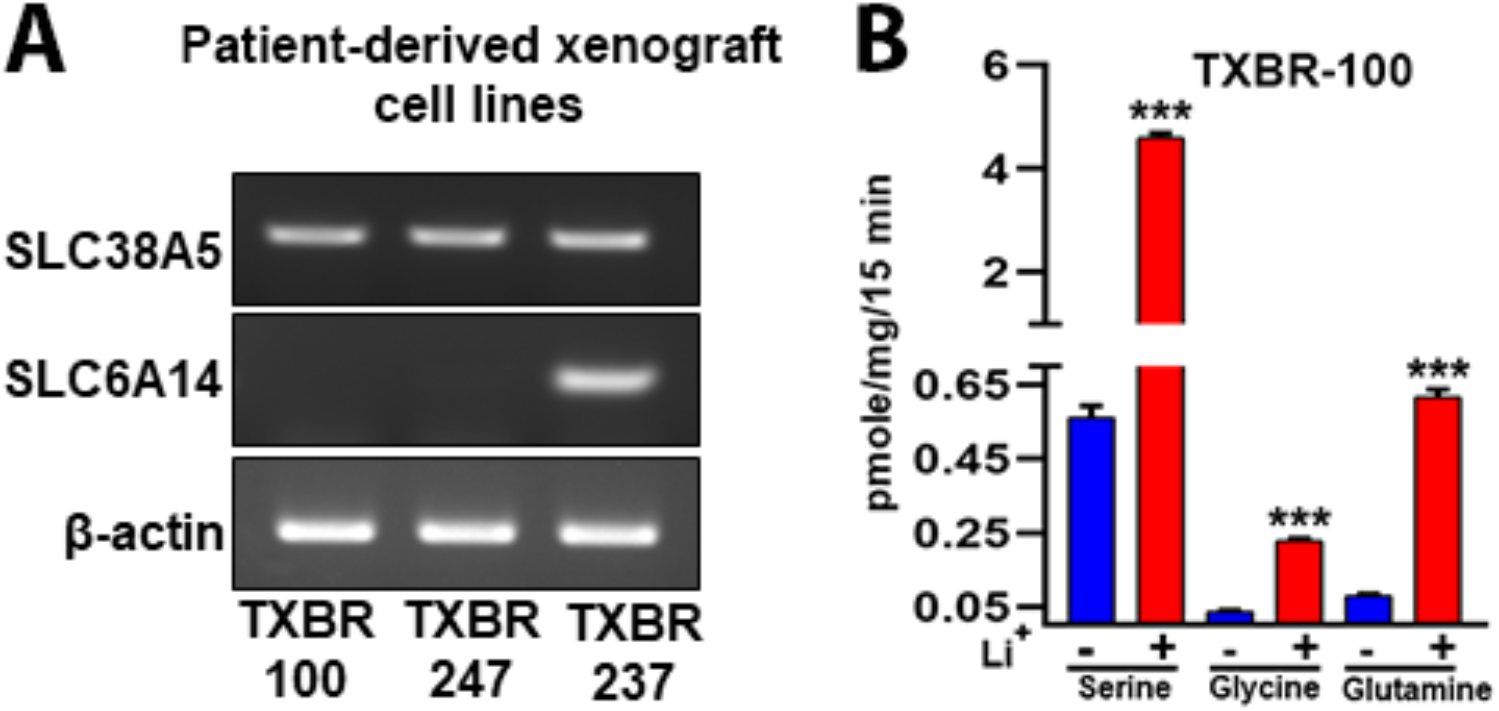
Expression and function of SLC38A5 in TNBC cell lines derived from patient-derived xenografts. (A) RT-PCR analysis of SLC38A5 and SLC6A14 mRNA expression in three cell lines. (B) Transport activity of SLC38A5 in the TNBC cell line TXBR-100 as monitored by the uptake of serine, glycine, and glutamine as the substrates for the transporter. The transport function of SLC38A5 was monitored in an uptake buffer (pH 8.5) containing 5 mM tryptophan to suppress the involvement of SLC7A5 in the uptake and by comparing the uptake in the presence and absence of Li^+^. The Li^+^-stimulatable uptake under these uptake conditions was taken as the transport activity specific for SLC38A5. ***, p<0.001 compared to uptake in the absence of Li^+^.

### Differential expression of Slc38A5 and Slc6a14 in three different mouse models of spontaneous breast cancer

Transgenic mouse lines are available for studies of spontaneously occurring breast tumors of different major subclasses: MMTV-Neu transgenic mouse line develops HER-2 positive breast cancer, MMTV-HRAS transgenic mouse line develops oncogene (HRAS)-driven breast cancer, and MMTV-PyMT mouse line develops ER+ breast cancer initially which then transforms into ER-negative breast cancer [26]. A similar change in terms of the ER status occurs also in ~30% of breast cancer in humans [27]. Normal virgin mammary tissue expresses neither Slc6a14 nor Slc38a5 (Fig. 4). We found that Neu-driven tumors and HRAS-driven tumors express only Slc6a14. Slc38a5 expression was completely absent in these two subclasses. In contrast, in tumors driven by polyoma middle T tumor antigen (PyMT), the expression of both Slc6a14 and Slc38a5 was clearly evident. We also monitored the expression of the other members of the SLC38 family and of the other amino acid transporters whose expression has been shown to be upregulated in breast cancer [6]. Slc38a1 and Slc38a2 are expressed at low levels in normal mammary tissue, and their expression is upregulated in all three subclasses of breast cancer. The expression of Slc38a1 is slightly increased in PyMT-driven breast cancer than in Neu- and HRAS-driven breast cancers. In contrast, there is no difference in the expression of Slc38a2 among the three subclasses. Slc1a5 shows an expression pattern similar to that of Slc38a1; it is expressed at low levels in normal mammary tissue and it is upregulated in all three subclasses of breast cancer. PyMT-driven breast cancer does show significantly higher expression of this transporter than in other breast cancer subtypes. Slc7a5 is expressed at negligible levels in normal mammary tissue and its expression is upregulated in breast cancer; however, there is no difference in expression among the three subclasses of breast cancer.

**Fig. 4.**
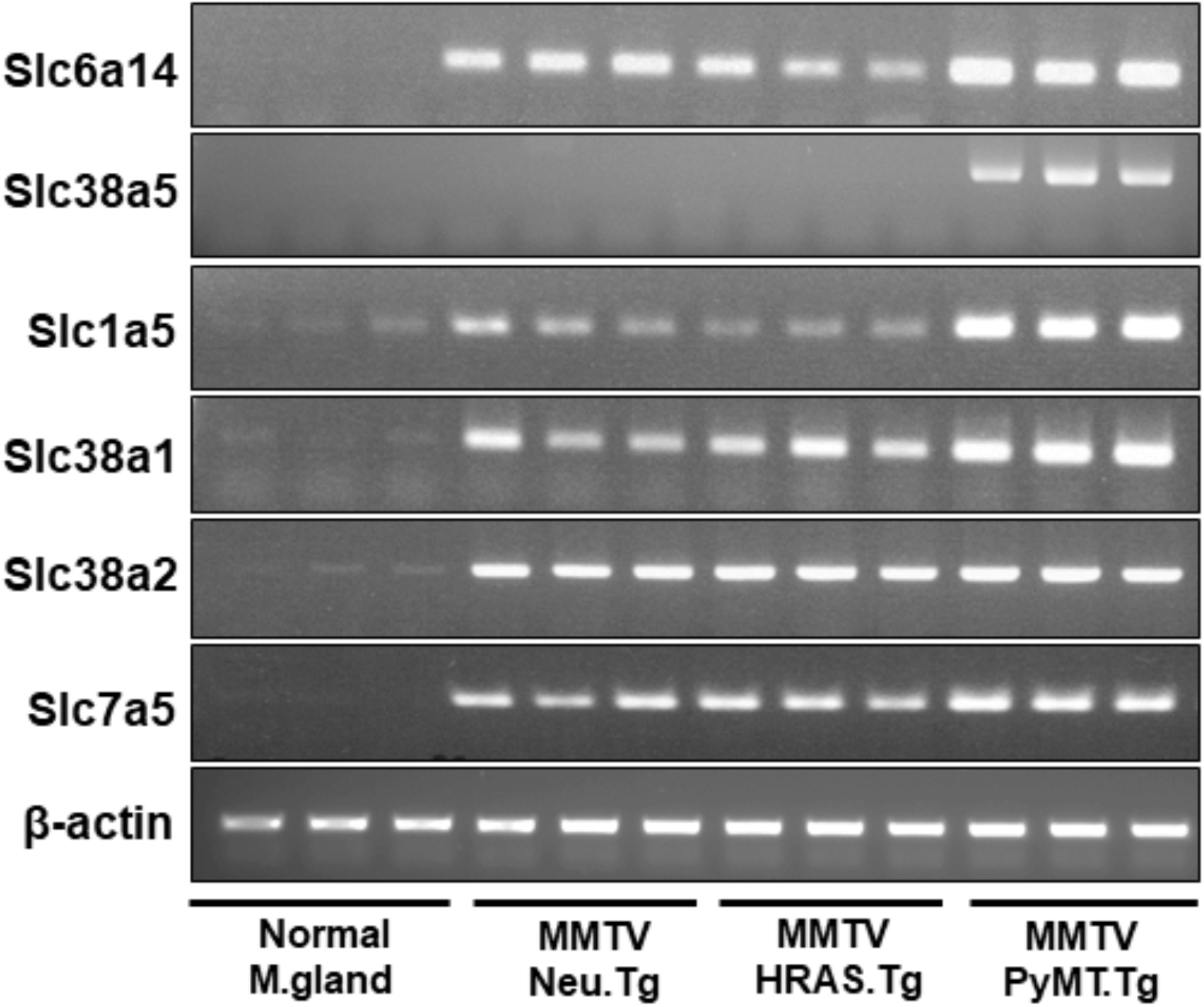
Differential expression of Slc38a5 in mouse models of spontaneous mammary tumors representative of different major subtypes. Tumors were collected from MMTV-Neu, MMTV-HRAS, and MMTV-PyMT female mice for RNA preparation. Mice used the study did not go through breeding or pregnancy. Accordingly, virgin adult female wild type mice were used to collect mammary gland as the control. Since the age at which spontaneous tumors develop in the three mouse lines varies markedly (2-3 months of age for MMTV-PyMT mice whereas 8-10 months for the other two lines), we did not use age-matched wild type mice as controls. Instead, we used 3-month-old adult mice for this purpose. RNA was used for RT-PCR to monitor the expression pattern of several amino acid transporters that might be of relevance to tumor growth.

### Functional features of Li^+^-coupled serine uptake in TNBC cell lines MB231 and TXBR-100

SLC38A5 is not only Li^+^-tolerant but also selective for specific amino acids and its ability to mediate the influx of amino acids into cells is coupled to an outwardly directed H^+^ gradient across the plasma membrane. To confirm that the TNBC cells do express this particular transporter at the functional level, we studied the features of Li^+^-dependent serine uptake in MB231 and TXBR-100 cells. First, we examined the substrate selectivity by monitoring the ability of various amino acids to compete with [^3^H]-serine (15 μM) for the uptake process. At a concentration of 5 mM, unlabeled serine caused 60-70% inhibition of [^3^H]-serine uptake (Fig. 5). The uptake was also inhibited by glutamine, histidine, glycine, and methionine. But, branched chain amino acids (valine, isoleucine, and leucine), aromatic amino acids (phenylalanine), anionic amino acids (aspartate and glutamate) and cationic amino acids (arginine and lysine) had no significant effect. Analysis of saturation kinetics indicated participation of a single saturable process with a Michaelis constant of 2.3 ± 0.2 mM for serine (data not shown). We then studied the effect of extracellular pH on SLC38A5-mediated serine uptake to determine the potential role of a H^+^ gradient in the transport process. The uptake was low at acidic pH, but it increased as the extracellular pH was increased (Fig. 6). As the extracellular pH is increased, the magnitude of the outwardly directed transmembrane H^+^ gradient across the plasma membrane increases proportionately. This suggests potential involvement of the outward-directed H^+^ gradient as a driving force for the uptake process.

**Fig. 5.**
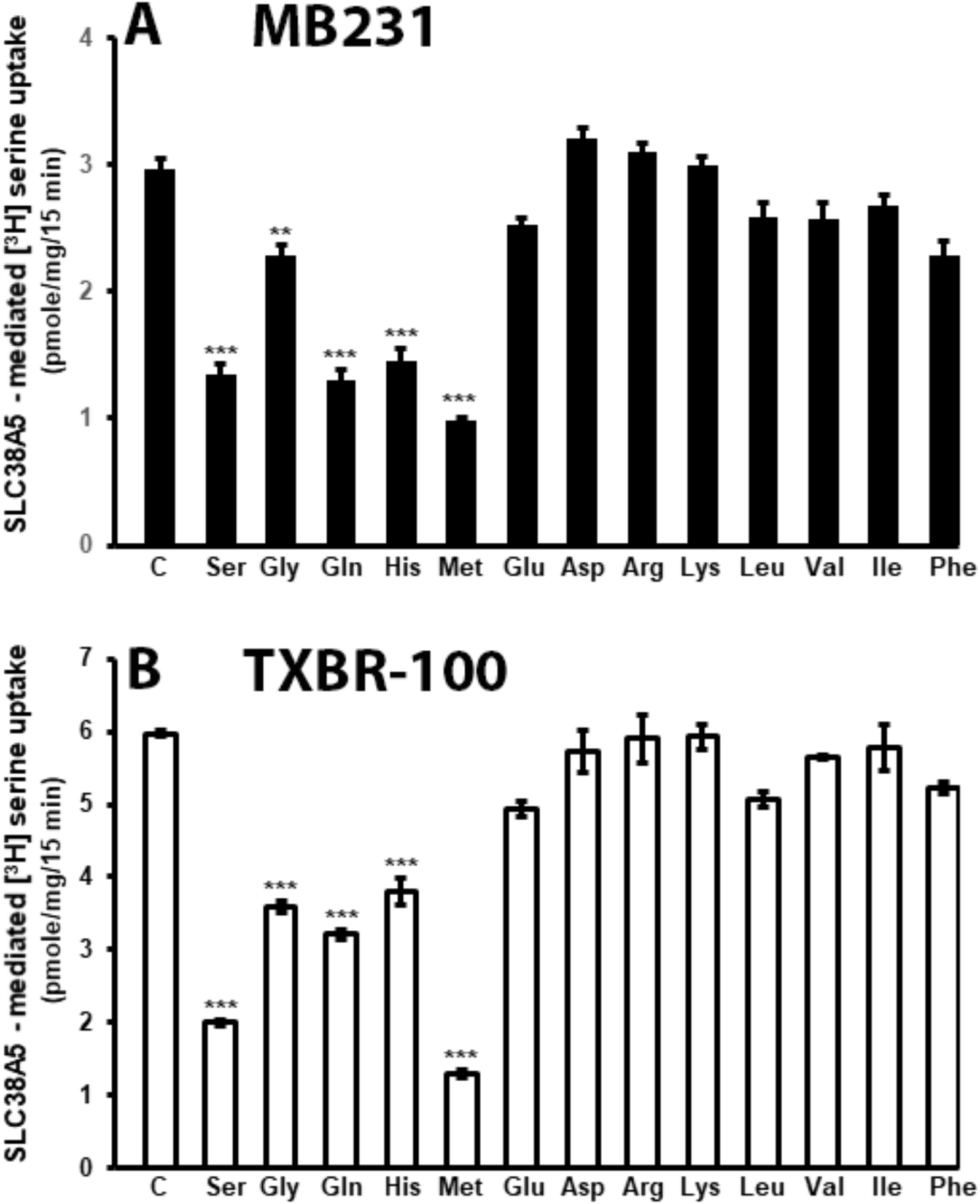
Substrate selectivity of Li^+^-coupled amino acid transport in TNBC cell lines. Uptake of [^3^H]-serine (15 μM) was measured in a LiCl-containing buffer, pH 8.5 (NaCl was replaced isosmotically with LiCl) in the absence or presence of various amino acids (5 mM). Uptake measured in the NMDGCl-buffer was subtracted from the uptake measured in the presence of LiCl (with or without the competing amino acids) to calculate the Li^+^-coupled uptake. In every case, the uptake buffer contained 5 mM tryptophan to suppress the contribution of serine uptake via SLC7A5. Data are means ± S. E. (n=6). Statistical analysis was done using the paired Student’s t test by comparing the uptake in the presence and absence of unlabeled amino acids. (A) MB231 cells; (B) TXBR-100 cells. ***, p<0.001 compared to control uptake (C).

**Fig. 6.**
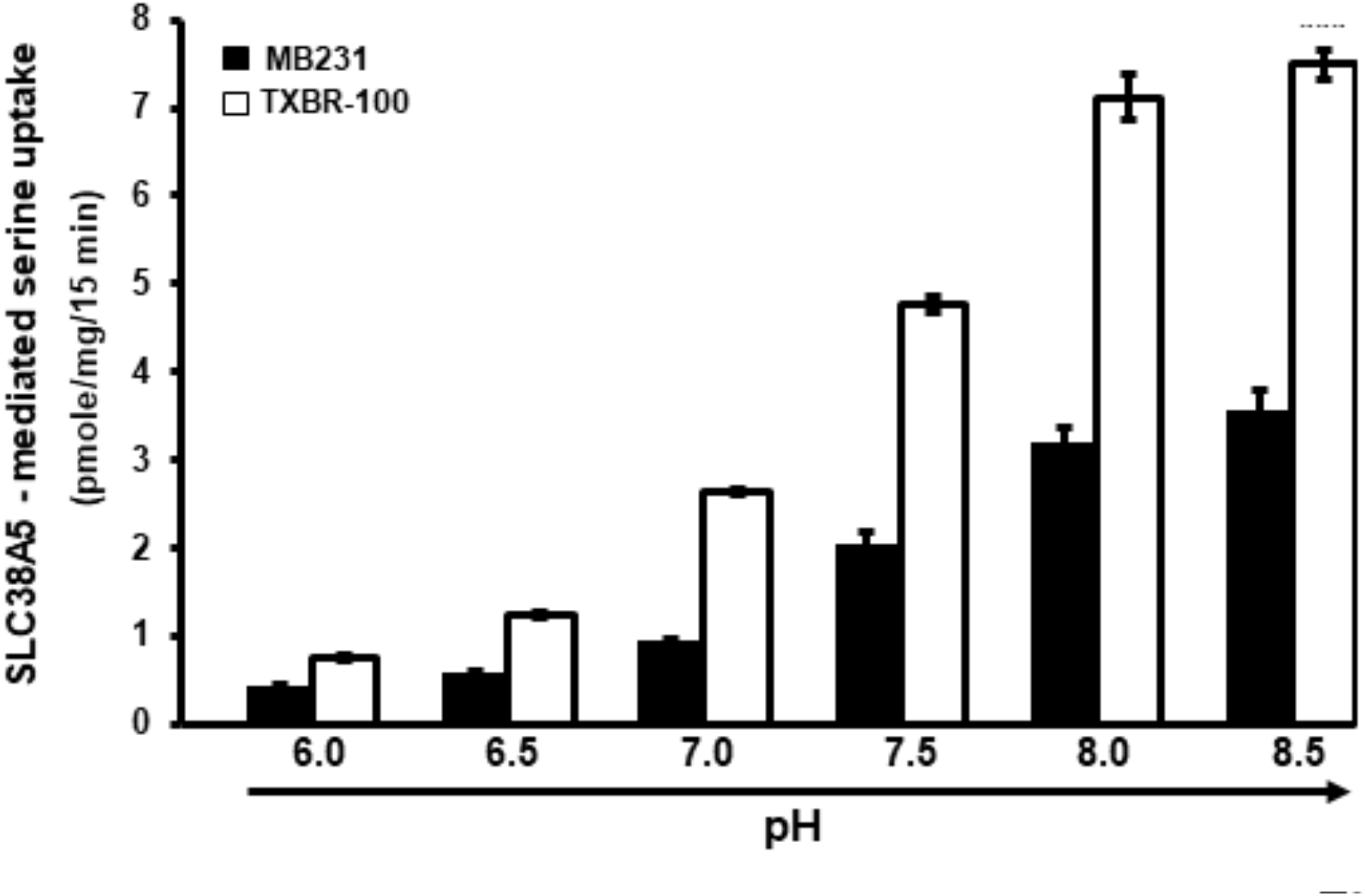
pH dependence of Li^+^-coupled [^3^H]-serine uptake (1 μM) in TNBC cell lines. Uptake buffer contained LiCl and 5 mM tryptophan. pH of the buffer was varied between 6.0 and 8.5. The pH 6.0 buffer was prepared using 25 mM Mes/Tris and the pH 8.5 buffer was prepared using 25 mM Tris/Hepes. The two buffers were mixed to prepare the buffers of other pH values. Serine uptake was measured in these buffers of different pH. Data are presented as means ± S.E. (n=6). ■, MB231 cells; □, TXBR-100 cells.

We also examined the impact of intracellular pH on the uptake process. For this, we used the NH_4_Cl prepulse method to cause intracellular acidification, another strategy to alter the transmembrane H^+^ gradient. We have used this method to demonstrate the energization of the peptide transporter PEPT1 [24] whose function is coupled to a transmembrane H^+^ gradient generated by Na^+^/H^+^ exchanger [28]. We found the uptake of serine (1 μM) in MB231 cells to be increased when intracellular pH was made acidic (control, 3.5 ± 0.2 pmoles/mg protein/15 min; NH_4_Cl prepulse, 4.7 ± 0.3 pmoles/mg protein/15 min; p<0.01), amounting to ~35% increase in uptake due to intracellular acidification. Serine uptake in TXBR-100 cells also responded to NH_4_Cl prepulse in a similar manner though the increase was a little bit lower (~25%) but still statistically significant (p<0.01).

### Functional coupling between SLC38A5-mediated amino acid uptake and macropinocytosis in TNBC cells

Based on the published reports that submembrane alkalinization on the cytoplasmic side of the plasma membrane caused by the activity of Na^+^/H^+^ exchanger promotes macropinocytosis [22, 23], we wondered whether the function of SLC38A5 as an amino acid-dependent Na^+^/H^+^ exchanger might also induce macropinocytosis. To investigate this, we monitored the process of macropinocytosis in TXBR-100 cells by the cellular entry of the fluorescent marker TMR-dextran. When studied in the presence of a NaCl-medium, the uptake of the fluorescent marker was markedly enhanced by serine, a substrate for SLC38A5 (Fig. 7A, B). In contrast, serine did not induce the uptake of TMR-dextran in a Na^+^-free medium. In fact, the uptake of the marker was drastically decreased in the absence of Na^+^. When Na^+^ is removed from the medium, serine uptake does not occur via SLC38A5 because its transport activity is obligatorily coupled to Na^+^ (or Li^+^) and at the same time Na^+^/H^+^ exchanger functions in the opposite direction. Under normal conditions with Na^+^ present in the extracellular medium, the Na^+^/H^+^ exchanger mediates the entry of Na^+^ into cells coupled to the efflux of H^+^ from the cells, thus causing intracellular alkalinization. But, when Na^+^ is absent in the extracellular medium, the exchanger mediates the efflux of Na^+^ from the cells coupled to the influx of H^+^ into cells, causing intracellular acidification. Since macropinocytosis is promoted by alkalinization on the cytoplasmic surface of the plasma membrane, which alters actin polymerization and hence impacts formation of macropinosomes, the decrease in the cellular uptake of the fluorescent marker for macropinocytosis is expected in a Na^+^-free extracellular medium.

**Fig. 7.**
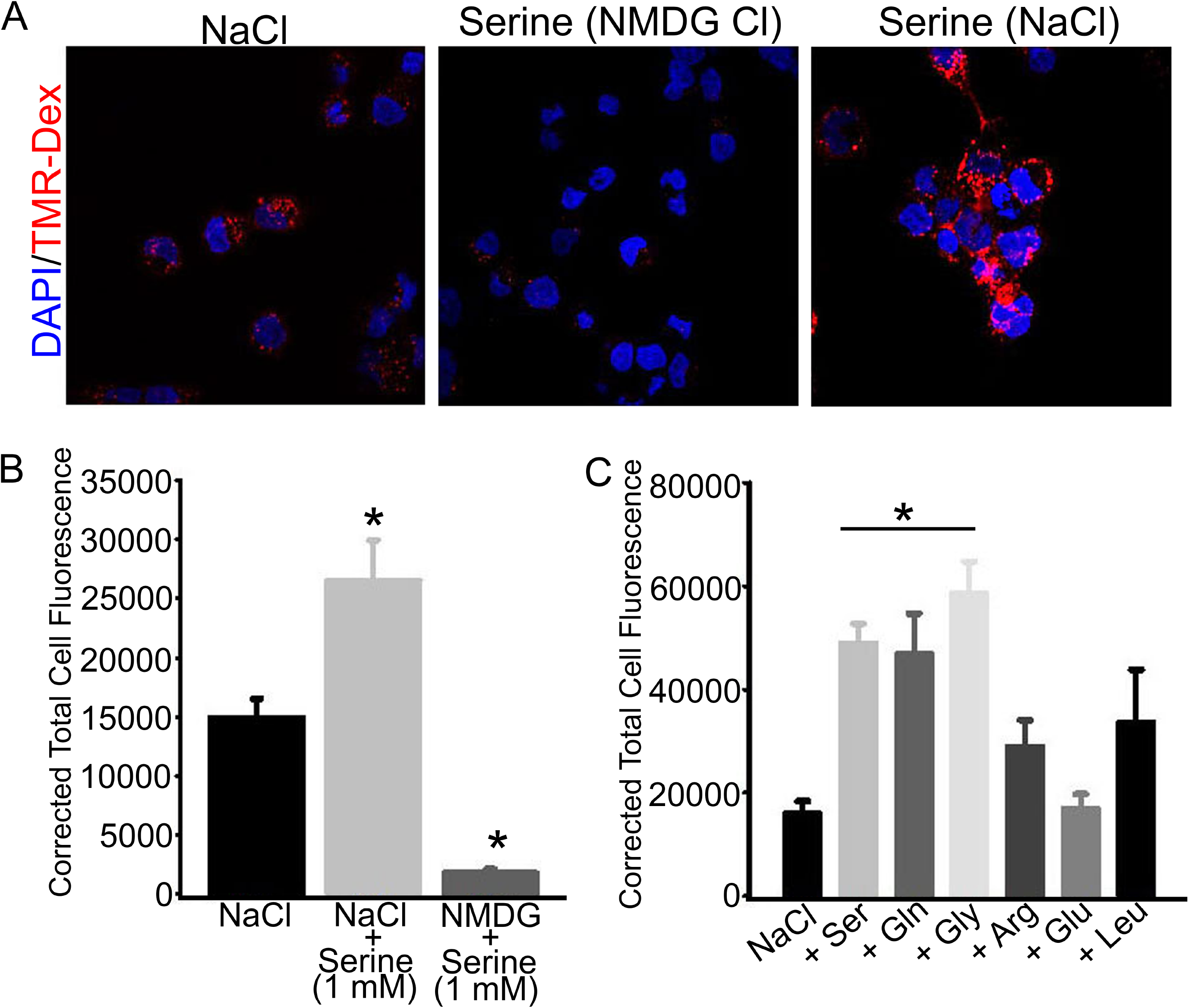
Promotion of macropinocytosis by SLC38A5-mediated transport function in the TNBC cell line TXBR-100. (A) Cells were incubated in buffers (pH 7.5) containing NaCl (Na^+^-buffer) or NMDGCl (Na^+^-free buffer) with and without serine (1 mM). All buffers contained TMR-dextran (100 μg/ml). The incubation was for 15 min at 37 °C. The cells were then fixed with 4% paraformaldehyde for 5 min, washed with phosphate-buffered saline, and mounted using Prolong diamond with DAPI. The images of TMR-dextran uptake (red) and DAPI (blue) were taken using confocal microscopy with a 60X objective and analyzed using NIS software. The images represent a maximum projection intensity derived from a Z-stack. (B) Quantification of the fluorescence signals, measured as the Corrected Total Cell Fluorescence using Image J software. ANOVA followed by Dunn’s test was used to determine the statistical significance among the groups. Data represent means ± S.E. *, p< 0.05. (C) Amino acid selectivity for the promotion of TMR-dextran uptake. Each amino acid was used at a concentration of 1 mM. Fluorescence signals were quantified and statistical significance analyzed as described above. *, p< 0.05.

If SLC38A5-mediated uptake of amino acids with its concomitant efflux of H^+^ was responsible for the serine-dependent increase in macropinocytosis in the NaCl-containing medium, this effect is expected to be restricted solely to the amino acid substrates of the transporter. To determine if this is true, we examined the influence of various amino acids on the uptake of the fluorescent marker. We found serine, glutamine, and glycine to be able to promote macropinocytosis (Fig. 7C). In contrast, arginine, glutamate, and leucine, which are not substrates for SLC38A5, failed to have any significant effect.

Amiloride and its derivatives are known inhibitors of macropinocytosis because of their ability to inhibit Na^+^/H^+^ exchanger [22, 23]. Since SLC38A5 also functions as a Na^+^/H^+^ exchanger but in an amino acid-dependent manner, we wondered whether the promotion of macropinocytosis caused by serine in a NaCl-containing extracellular medium was sensitive to inhibition by amiloride. We addressed this by examining the effect of ethylisopropyl amiloride (EIPA) on Na^+^/serine-induced increase in the uptake of the fluorescent marker (Fig. 8). EIPA blocked macropinocytosis almost completely under these conditions. EIPA inhibited the uptake of the fluorescent marker even in the absence of serine because it is an inhibitor of Na^+^/H^+^ exchanger. But, it was interesting that EIPA also blocked the serine-induced increase in the uptake of the marker, suggesting that this amiloride derivative might also be a blocker of the amino acid-dependent Na^+^/H^+^ activity of SLC38A5.

**Fig. 8.**
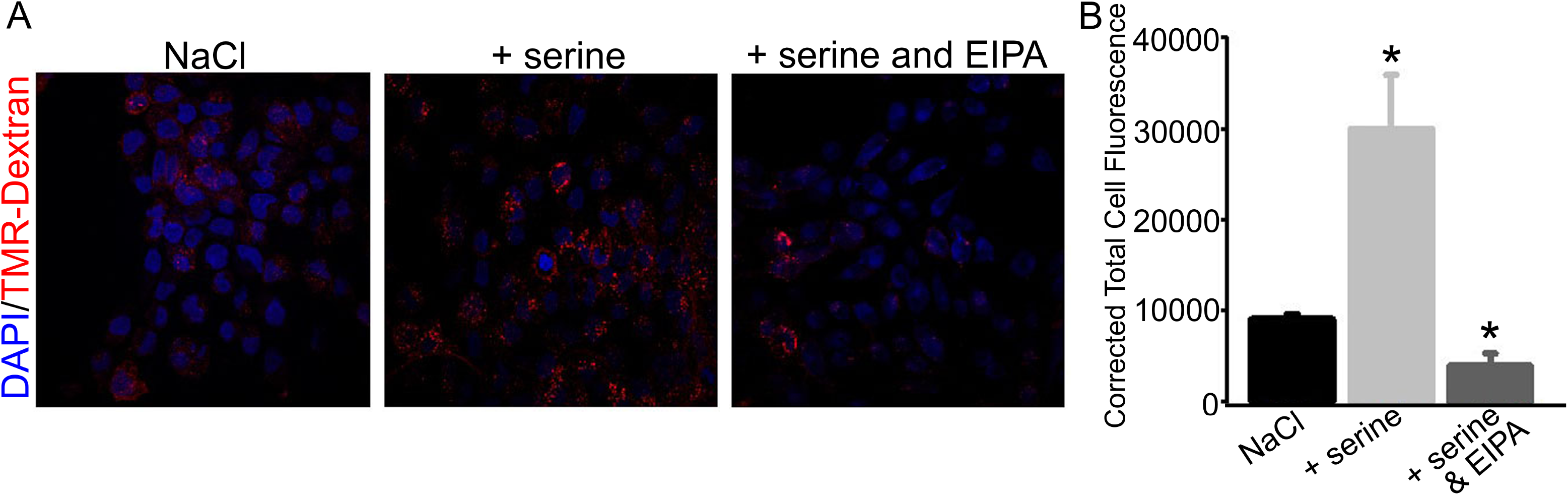
Blockade of SLC38A5-induced macropinocytosis by EIPA (ethylisopropyl amiloride). (A) Macropinocytosis was monitored using TMR-dextran in three different buffers (pH 7.5): NaCl, NaCl plus serine (1 mM), and NaCl plus serine (1 mM) and EIPA (100 μM). (B) Quantification of the fluorescence signals, and statistical significance among the groups. Data represent means ± S.E. *, p< 0.05.

To further confirm the role of SLC38A5 as a promoter of macropinocytosis, we employed experimental conditions in which serine uptake occurs solely via SLC38A5. This was done by performing the experiments in the presence of 5 mM tryptophan, which blocks the uptake of serine via the amino acid transporter SLC7A5. Another change in the experimental condition was the replacement of NaCl with LiCl, thus restricting the uptake of serine solely to SLC38A5 without the participation of any other serine uptake system that might be coupled to Na^+^. We found that, in the presence of tryptophan, serine promoted macropinocytosis both NaCl medium and LiCl medium (Supplemental Fig. S1A, B). We also monitored serine-induced macropinocytosis in MB231 cells that have robust SLC38A5 expression and activity and also in the ER+ breast cancer cell line ZR 75.1 that have no SLC38A5 expression and activity (Fig. 2). When the experiment was performed in a NaCl-containing medium, serine induced macropinocytosis in MB231 cells but not in ZR 75.1 cells (Supplemental Fig. S2).

Taken collectively, these experiments demonstrate that amino acid transport via SLC38A5 is coupled to promotion of macropinocytosis in TNBC cells and that the amino acid-dependent Na^+^/H^+^ exchange activity of the transporter underlies this phenomenon.

### Inhibition of SLC38A5 transport function by amiloride and its derivatives

Amiloride and its derivatives are widely used as inhibitors of Na^+^/H^+^ exchangers. Since SLC38A5 functions as an amino acid-dependent Na^+^/H^+^ exchanger and it also induces macropinocytosis as the classical Na^+^/H^+^ exchanger does, we were curious to find out if amilorides would interact with SLC38A5 and interfere with its transport function. For this, we evaluated the effects of amiloride and its derivatives on SLC38A5-mediated serine transport in MB231 cells. The uptake was measured in a LiCl-buffer, pH 8.5, containing 5 mM tryptophan, the conditions that allow us to specifically monitor the SLC38A5-specific serine uptake. All four amilorides examined in the study inhibited SLC38A5 (Fig. 9). The most effective amilorides were ethylisopropyl amiloride and hexamethylene amiloride. Harmaline is known to interfere with Na^+^-coupled transport systems [29]. We found this compound also to have the ability to inhibit the transport function of SLC38A5 (Fig. 9).

**Fig. 9.**
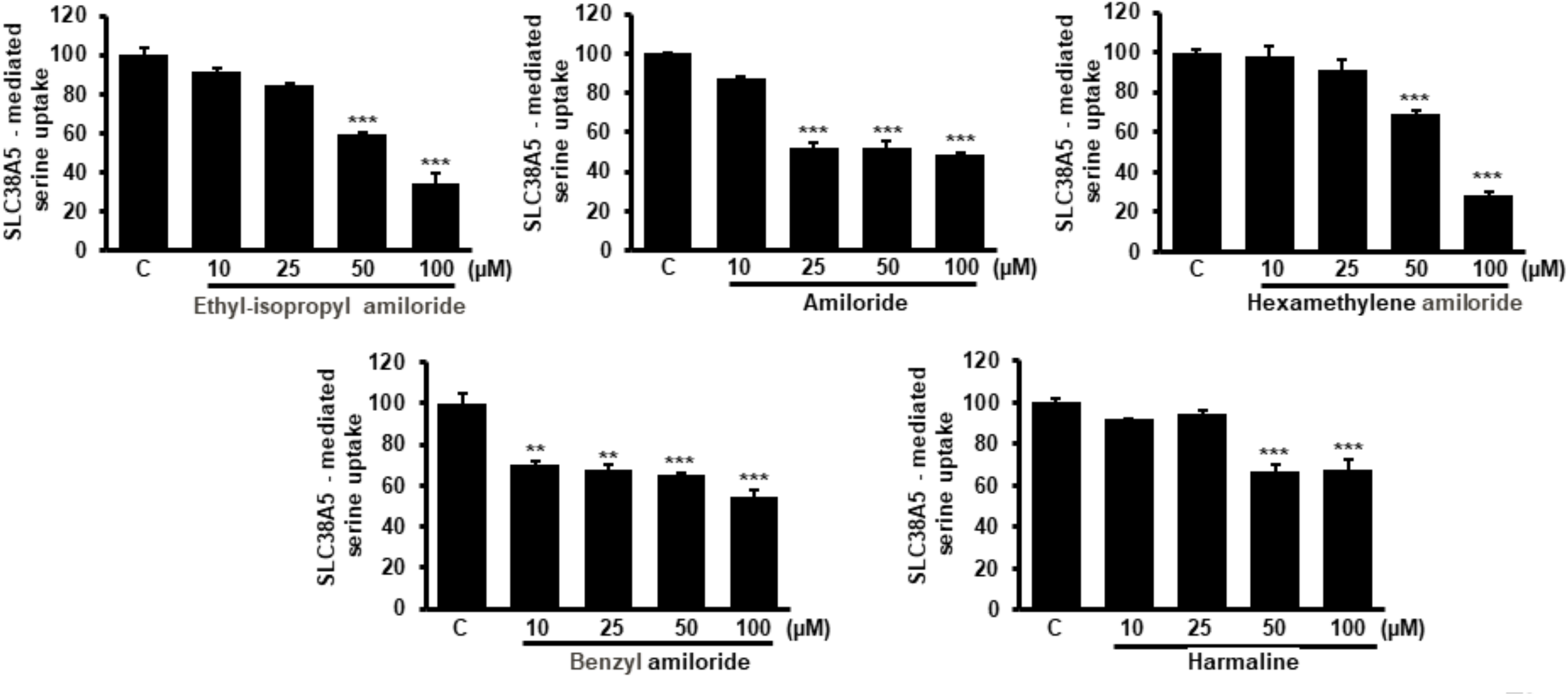
Inhibition of SLC38A5-mediated [^3^H]-serine uptake (1 μM) by amiloride and its derivatives. Serine uptake was measured for 15 min in LiCl-buffer (pH 8.5) containing 5 mM tryptophan. Amiloride and its derivatives were present during uptake at increasing concentrations. Control uptake (C) was taken as 100% in each experiment. Data represent means ± S.E. (n = 6). ANOVA followed by Dunn’s test was used to determine the statistical significance among the groups. **, p<0.01; ***, p<0.001.

### Differential expression of SLC38A5 and SLC38A2 in different subtypes of breast cancer

The present investigation focuses on the expression of SLC38A5 in breast cancer. There have been two recent reports on the expression of SLC38A2, another member of the SLC38 family, in breast cancer [16, 17]. SLC38A5 represents one of the subtypes of the amino acid transport system N, the other being SLC38A3; both function to mediate the influx of their amino acid substrates into cells coupled to Na^+^ symport and H^+^ antiport [19, 20, 30]. SLC38A2 is also a Na^+^-coupled amino acid transporter whose functional features resemble those of the amino acid transport system A [18], but there is no involvement of H^+^ in the transport mechanism associated with SLC38A2. SLC38A5 and SLC38A2 also differ in substrate specificity; the former possesses a much narrower amino acid selectivity whereas the latter is capable of transporting almost all neutral amino acids. We did not find any differential expression of SLC38A2 among the human breast cancer cells representative of ER+ or ER-negative breast cancer. A recently published report has shown that SLC38A2 expression is associated with oxidative stress resistance and poorer prognosis in TNBC [17]. Given that SLC38A5 as well as SLC38A2 might have relevance to breast cancer growth and progression, we analyzed the expression of these two transporters in breast cancer using the TCGA (The Cancer Genome Atlas) database. We found the expression of SLC38A5 to be significantly higher in primary breast tumor tissues than in normal breast tissue (Fig. S3A). The overexpression was evident in all three major subclasses of breast cancer (ER+ or luminal, HER2+, and basal or TNBC) (Fig. S3B). When the expression in ER+ and ER-negative breast cancer subtypes was analyzed separately, we found the expression to be much higher in ER-negative (TNBC) subtype than in ER+ (non-TNBC) subtype (Fig. S4). These observations corroborate the data from the present study. In contrast, the expression of SLC38A2 is reduced in primary breast cancer tissues compared to normal breast tissue (Fig. S5A). The decrease in expression was seen in all three major classes of breast cancer across the board; the most prominent downregulation however was in TNBC (basal subtype) (Fig. S5B). Similarly, the expression of SLC38A3 was also found to be decreased in ER+, HER2+, and TNBC compared to normal mammary tissue (TCGA database).

## Discussion

The findings of the present study can be summarized as follows: (i) SLC6A14 and SLC38A5 are expressed differentially in ER+ breast cancer and TNBC; SLC6A14 is upregulated in ER+ breast cancer whereas SLC38A5 is upregulated in TNBC; (ii) the differential expression of these two transporters in different subtypes of breast cancer is evident also in cell lines representative of the two subtypes; (iii) there is no difference in the expression of other membranes of the SLC38 family between ER+ breast cancer and TNBC; (iv) the selective expression of SLC38A5 in TNBC is also observed in mouse models of spontaneous breast cancer for the different subtypes; (v) the transport function of SLC38A5 is demonstrable both in conventional TNBC cell lines and in patient-derived xenograft cell lines; (vi) SLC38A5 promotes macropinocytosis in TNBC cells; SLC38A5 is known to function as an amino acid-dependent Na^+^/H^+^ exchanger, a unique functional feature of the transporter, which seems to underlie the ability of SLC38A5 to promote macropinocytosis; (vii) the small molecule inhibitors of the classical Na^+^/H^+^ exchangers (members of the SLC9 family) also inhibit the transport function of SLC38A5, though not with the same potency.

The Na^+^/Cl^-^ -coupled broad-selective amino acid transporter SLC6A14 is expressed ER+ breast cancer cell lines but not in TNBC cell lines [25]. This is true also in primary breast tumor tissues [25]. SLC6A14 is a direct target for estrogen/ER, thus providing the molecular basis for its upregulation in ER+ breast cancer [25]. Furthermore, deletion of Slc6a14 in mice interferes with the growth of spontaneous mammary tumors [31]. These studies raised the question as to the identity of the transporter that is responsible for satisfying the amino acid needs in TNBC. Previous reports have shown that the amino acid transporters SLC1A5, SLC7A5, and SLC7A11 are upregulated in TNBC [5–7]. But, the functional features of these three transporters are not ideally suited to meet the increased demands for amino acids in cancer cells [6]. SLC1A5 and SLC7A5 are obligatory amino acid exchangers, meaning that when these transporters mediate the influx of one of their amino acid substrates into cells, one of their other amino acid substrates is removed from the cells. Such a mechanism may be suitable for intracellular accumulation of specific amino acids for specialized intracellular functions as has been shown for leucine and its associated mTORC1 signaling [32]. But, this mechanism is not ideal for the general supply of amino acids to promote amino acid nutrition. SLC7A11 is also an exchanger with selectivity towards only cystine and glutamate; it transports cystine into cells coupled to the efflux of glutamate [33]. As such, this transporter is ideal to supply cysteine for glutathione synthesis and promote antioxidant machinery in cancer cells but not to meet the increased demands for amino acids. Therefore, we envisaged that there should be some other transporter(s) which is/are upregulated in TNBC to supply amino acids in support of various metabolic pathways that are reprogrammed in cancer. We thought that SLC38A5 is an excellent candidate for TNBC because of its unique functional features [14,19,20]. It transports glutamine, glycine, serine, and methionine, coupled to the transmembrane Na^+^ gradient [19,20]. These amino acids are expected to maintain two important metabolic pathways that are accelerated in cancer, namely glutaminolysis associated with ATP production and lipid synthesis, and one-carbon metabolism associated with nucleotide synthesis and epigenetic control of gene expression [14]. The transporter is also coupled to H^+^ efflux, meaning that when amino acids are transported into cells via SLC38A5, the process is accompanied with the removal of H^+^ from the cells, thus leading to intracellular alkalinization [20]. Cancer cells would benefit from this process because intracellular alkalinization promotes mitogenesis [34] and macropinocytosis [22,23]. But, there have been no published reports investigating the expression of this amino acid transporter in breast cancer or any other cancers.

Whenever amino acid transporters in cancer are investigated, the sole focus is on the role of these transporters in the provision of amino acids to cancer cells. The present investigation departs from this norm; it focuses on the function of an amino acid transporter, which also possesses Na^+^/H^+^ exchange activity, and its relation to the endocytic process known as macropinocytosis. Since macropinocytosis facilitates the cellular entry of macromolecules, including proteins, present in the extracellular fluid, SLC38A5 contributes to amino acid nutrition in TNBC by two totally independent mechanisms, firstly by the traditional delivery of amino acids via the function of SLC38A5 as an amino acid transporter, and secondly by the promotion of macropinocytosis via the function of SLC38A5 as an amino acid-dependent Na^+^/H^+^ exchanger. This process of macropinocytosis delivers extracellular proteins to lysosomes with subsequent degradation of the proteins into amino acids for utilization by the cells.

The functional feature that is fundamental to the ability of SLC38A5 to promote macropinocytosis is its Na^+^/H^+^ exchange activity. SLC38A3 and SLC38A5 were once thought to be electrogenic transporters based on the inward currents induced by amino acid substrates in the presence of Na^+^ in frog oocytes that expressed these transporters heterologously [30]. But later studies showed that despite the fact that amino acids induce inward currents, this phenomenon is not related to amino acid transport per se [35, 36]. Amino acid transfer into cells via SLC38A5 is electroneutral, with one Na^+^ going into the cells coupled to the efflux of H^+^. The inward current, seen when SLC38A5-expressing oocytes are exposed to amino acid substrates of the transporter, has nothing to do with amino acid transfer but is related to amino acid-induced cation conductance. It is this cation conductance, not related to the entry of amino acids, that contributes to the inward currents. Whether this feature has any significance in vivo or whether it is a phenomenon that occurs only in the oocyte expression system needs to be investigated. The Na^+^/H^+^ exchange feature is unique only to SLC38A3 and SLC38A5. Schneider et al [36] have discovered that SLC38A3 does not function only as an amino acid-dependent Na^+^/H^+^ exchanger; the transporter possesses Na^+^/H^+^ exchange activity that is not coupled to amino acid transport. In fact, the uncoupled Na^+^/H^+^ activity may be higher in magnitude than the activity coupled to amino acid transport. It is possible that a similar phenomenon also occurs with SLC38A5, which might have direct relevance to the transporter’s ability to induce macropinocytosis. However, in the present study, we focused solely on the amino acid-dependent Na^+^/H^+^ exchange. There are about three dozen amino acid transporters in mammalian cells, and to the best of our knowledge, no amino acid transporter other than SLC38A3 and SLC38A5 possesses this interesting feature of Na^+^/H^+^ exchange. This is an important characteristic with direct relevance to cancer. When amino acids enter cancer cells via SLC38A5, it promotes the efflux of H^+^, which might contribute to the maintenance of intracellular pH. Cancer cells have an increased risk for intracellular acidification because of production of lactic acid in aerobic glycolysis [3]. Here we have unraveled another interesting and important relationship between SLC38A5 and cancer cell biology. The amino acid-dependent Na^+^/H^+^ exchange activity of SLC38A5 promotes macropinocytosis, which may contribute significantly to amino acid nutrition in cancer cells. The fact that this transporter is selectively upregulated in TNBC underscores its importance as a tumor promoter in TNBC. Even though SLC38A3 exhibits functional features that are almost identical to those of SLC38A5 [7], the expression of the former is actually decreased in breast cancer, and the decrease is independent of the hormone receptor status of the tumor (TCGA database). This suggests that SLC38A5 might have potential as an actionable drug target for TNBC. At present, there are no targeted therapies available for TNBC and SLC38A5 might fill this void and offer a unique therapeutic target.

The pH dependence of SLC38A5 deserves some discussion with regard to the relevance of this transporter to cancer. Since the transporter function involves the movement of Na^+^ and amino acids in one direction coupled to the movement of H^+^ in the opposite direction, the influx of amino acids into cells in the presence of an inwardly directed Na^+^ gradient via the transporter is stimulated by an alkaline pH in the extracellular medium. The influx of amino acids is suppressed when the extracellular medium is acidic. In solid tumors such as the breast cancer, the tumor microenvironment is acidic, the conditions under which the influx of amino acids into cancer cells via the transporter would be low. It is possible that SLC38A5 plays a role in the transport of amino acids into cancer cells during initial stages of carcinogenesis when the extracellular pH is not acidic. This is also the period when the function of the transporter would be associated with the promotion of macropinocytosis, thus providing additional mechanisms for the cancer cells to acquire nutrients. As the pH in the extracellular medium turns more acidic with time during tumor growth, the transporter might assume different functions. For example, the transporter might work in the reverse direction by facilitating the efflux of Na^+^ and amino acids from the cancer cells in exchange for extracellular H^+^. In normal physiology, this potential for bidirectional function of SLC38A3 and SLC38A5 is important in the glutamine-glutamate cycle that occurs in the brain between glutamatergic neurons and the surrounding astrocytes [37, 38]. In this process, neurons release glutamate by exocytosis upon activation to elicit neurotransmission; then, the released glutamate is taken up by astrocytes via glutamate transporters to be converted into glutamine. This glutamine is then released into the extracellular fluid via SLC38A3 and SLC38A5, which is then taken up by neurons via SLC38A1 and SLC38A2 for subsequent conversion into glutamate for reuse as the neurotransmitter. In this scenario, SLC38A3 and SLC38A5 function in the reverse mode for the release of glutamine from the cells. SLC38A5 could carry out a similar function in solid tumors at advanced stages because of the acidic tumor microenvironment. Considering that glutamine, glycine, and cysteine are good substrates for SLC38A5, efflux of these amino acids via the transporter might result in decreased cellular levels of the antioxidant glutathione (g-glutamyl-cysteinyl-glycine) in advanced tumors. This could potentially be relevant to sensitivity and responsiveness of the tumors to selective chemotherapeutic agents whose clearance from the cells is dependent on the mercapturic pathway involving glutathione [39]. Tumors with high expression of SLC38A5 might be more sensitive to such drugs than tumors with low expression of SLC38A5; this is however only a hypothesis at present, but it is a concept that deserves further investigation in the future.

## Data Availability Statement

All supporting data in relation to the studies reported here are provided in this manuscript.

