## Supplementary Materials for "Expression and function of SLC38A5, an amino acid-coupled Na^+^/H^+^ exchanger, in triple-negative breast cancer and its relevance to macropinocytosis"

### Supplementary Material

**Supplementary Table 1.** PCR primer sequences

Human-specific primers

| Gene | Forward Primer | Reverse Primer |
|---|---|---|
| **SLC6A14** | ATC GTC TGG CAA GGT GGT AT | TGA GTG GCA GCA TCT TTC CAT |
| **SLC38A5** | GTT GGG GCC ATG TCC AGT TA | AGT GTT TCA TGA GGG CGA GG |
| **SLC1A5** | GAG ACT CCA AGG GGC TCG C | CAC AAG CAG GTT GGC TCG AAG |
| **SLC38A1** | TTT GGA GTC GTA GGA GTT ACA TCT | TGG AAA CTG GAG GAA GAG AAA GA |
| **SLC38A2** | GCA GTG GAA TCC TTG GGC TT | ATA AAG ACC CTC CTT CAT TGG CA |
| **SLC7A5** | CGC TCT TCC CCA CCT GC | GAC ACA TCA CCC TTC CCG AT |
| **GAPDH** | CCA CTC CTC CAC CTT TGA C | ACC CTG TTG CTG TAG CCA |
| **HPRT** | GCG TCG TGA TTA GCG ATG ATG AAC | CCT CCC ATC TCC TTC ATG ACA TCT |

Mouse-specific primers

| Gene | Forward Primer | Reverse Primer |
|---|---|---|
| **Slc6a14** | CCA ATG GCG GAG GTG CTT TC | ATC AGG ACC ATC GTG ATT CCC |
| **Slc38a5** | CAC AAC GTT GGG GCT ATG TC | TGC TTG TGT ACC CCA GGT AG |
| **Slc1a5** | TTC CCC TCC AAT CTG GTG TCT | CCA CCT CAC AGA GAA GCT GGA C |
| **Slc38a1** | ACG CGT GCA CAC CAA AGT AT | AAA GAT GGC CGT CAG GAA GT |
| **Slc38a2** | TTG CAG GCC ACG CTA TTT CA | AGC ACA GCC AAT CGG ACA ACA A |
| **Slc7a5** | GCT GAC GAA CCT GGC CTA TT | TAG TTC CCG AAG TCC ACA GC |
| **β-actin** | CTG GCA CCA CAC CTT CTA | GGG CAC AGT GTG GGT GAC |

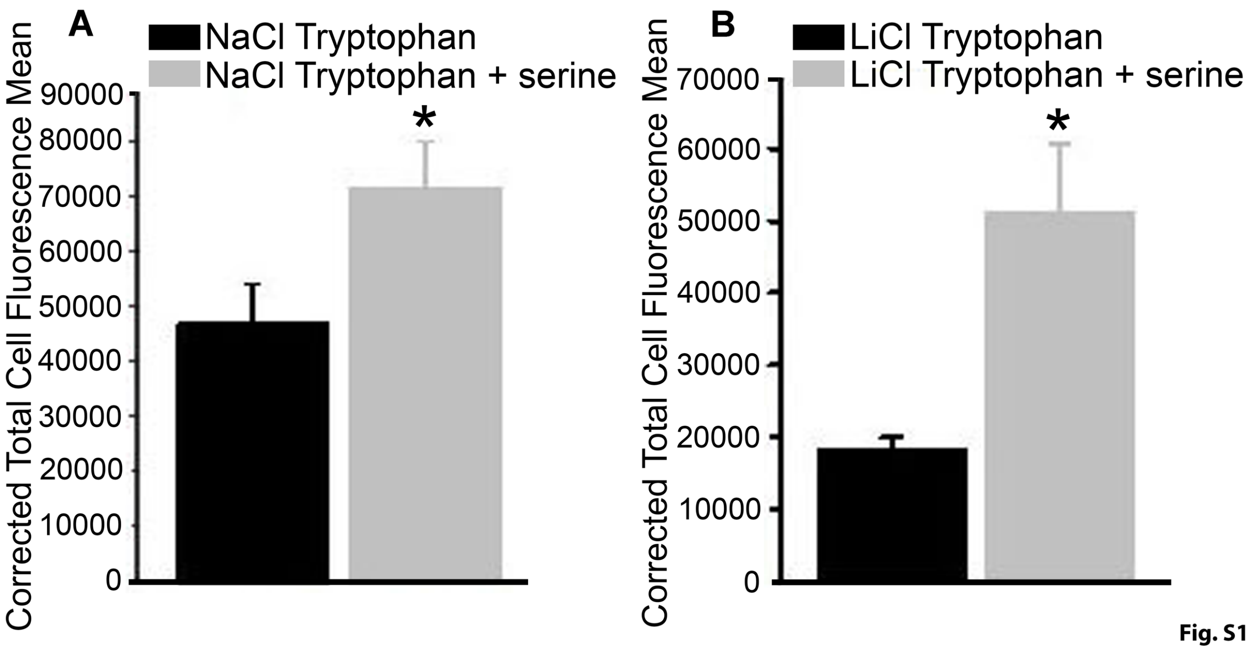


**Fig. S1.** Serine-induced macropinocytosis in the presence of $Na^+$ (A) or $Li^+$ (B) in the TNBC cell line TXBR-100. Macropinocytosis was monitored by the uptake of TMR-dextran by measuring intracellular fluorescence. Cells were incubated with TMR-dextran in NaCl-buffer (pH 7.5) or LiCl-buffer (pH 7.5) containing 5 mM tryptophan with and without serine (1 mM). Fluorescence signals were quantified and reported as Corrected Total Cell Flourescence (means ± S. E.).

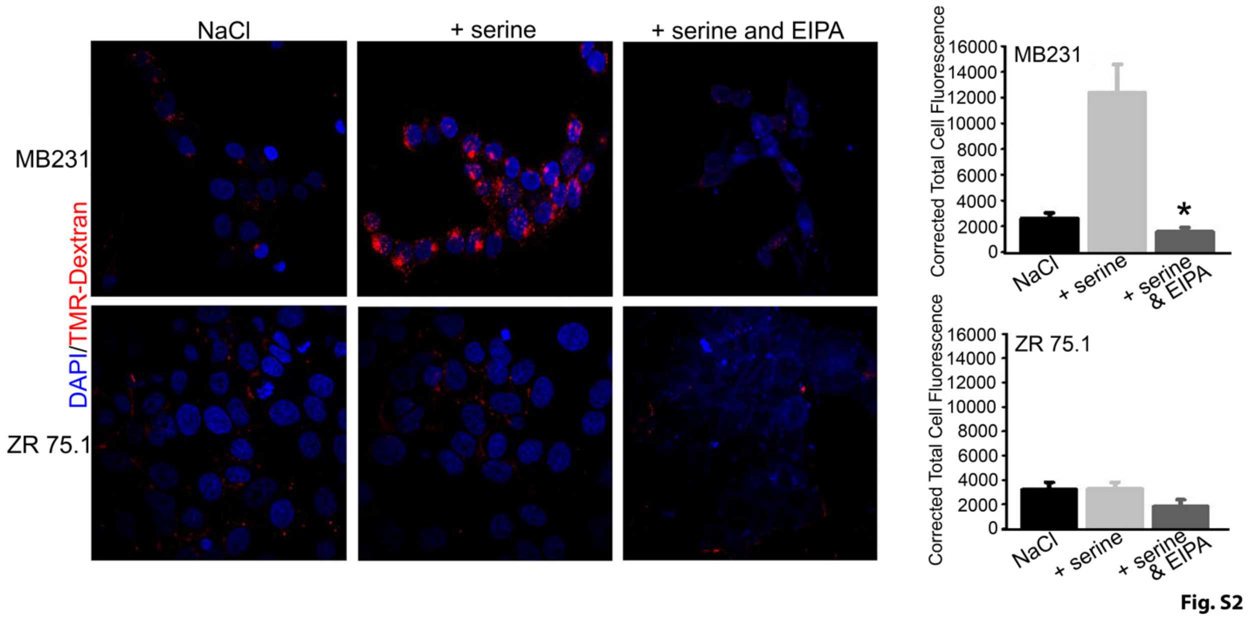


**Fig. S2.** Differential influence of serine on macropinocytosis in the TNBC cell line MB231 and the ER+ cell line ZR 75.1. Macropinocytosis was monitored using TMR-dextran in NaCl-buffer with and without serine (1 mM) and with and without EIPA (100 µM). The fluorescence signals were quantified and reported as Corrected Total Cell Fluorescence (means ± S.E.).

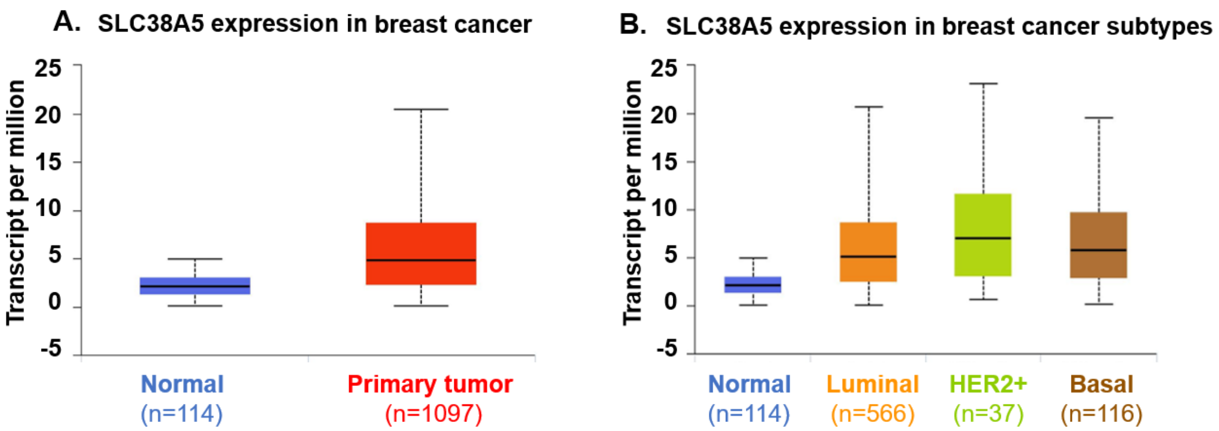


| Comparision | Statistical Significance |
|---|---|
| Normal vs Primary | 1.6E-12 |

| Comparision | Statistical Significance |
|---|---|
| Normal vs Luminal | <E-12 |
| Normal vs HER2+ | 2.95E-07 |
| Normal vs Basal | 3.87E-07 |

**Fig. S3**

**Fig. S3.** Expression pattern for SLC38A5 in breast cancer. (A) The TCGA database was used to analyze the expression level of SLC38A5 mRNA in primary breast tumor tissues and in normal mammary gland tissue. (B) The same database was used to analyze the expression pattern for SLC38A5 mRNA in normal mammary gland and in breast cancer of the three major breast cancer subtypes. The numbers in parentheses represent the number of cases in each category.

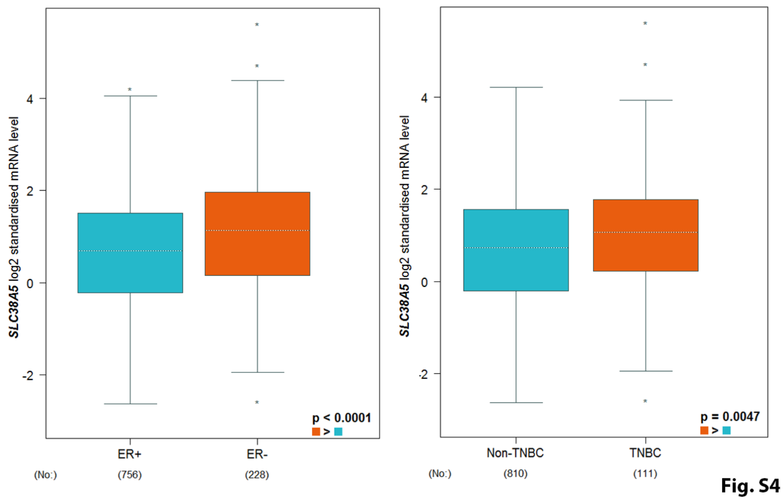

Fig. S4

**Fig. S4.** Relative expression of SLC38A5 mRNA in ER+ versus ER-negative breast cancer, and in TNBC versus non-TNBC. The information in the TCGA database was accessed and reformatted. The numbers in parentheses denote the number of cases in each category.

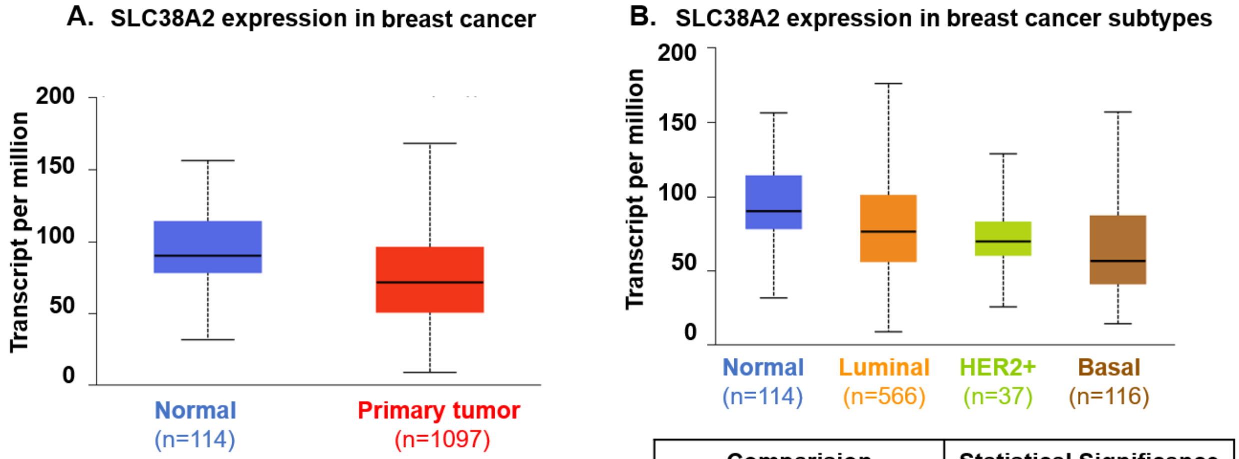


| Comparision | Statistical Significance |
|---|---|
| Normal vs Primary | 1.1E-6 |

| Comparision | Statistical Significance |
|---|---|
| Normal vs Luminal | 9.4E-04 |
| Normal vs HER2+ | 5.2E-03 |
| Normal vs Basal | 1.5E-08 |

**Fig. S5**

**Fig. S5.** (A) Expression pattern for SLC38A2 mRNA in primary breast cancer tissues and in non-involved normal tissues from the mammary gland. (B) Expression pattern of SLC38A5 in normal mammary gland and in the three major subtypes of breast cancer. The numbers in parentheses represent the number of cases in each category.
